## Supplementary Figures for "A neural network model that generates salt concentration memory-dependent chemotaxis in *Caenorhabditis elegans*"

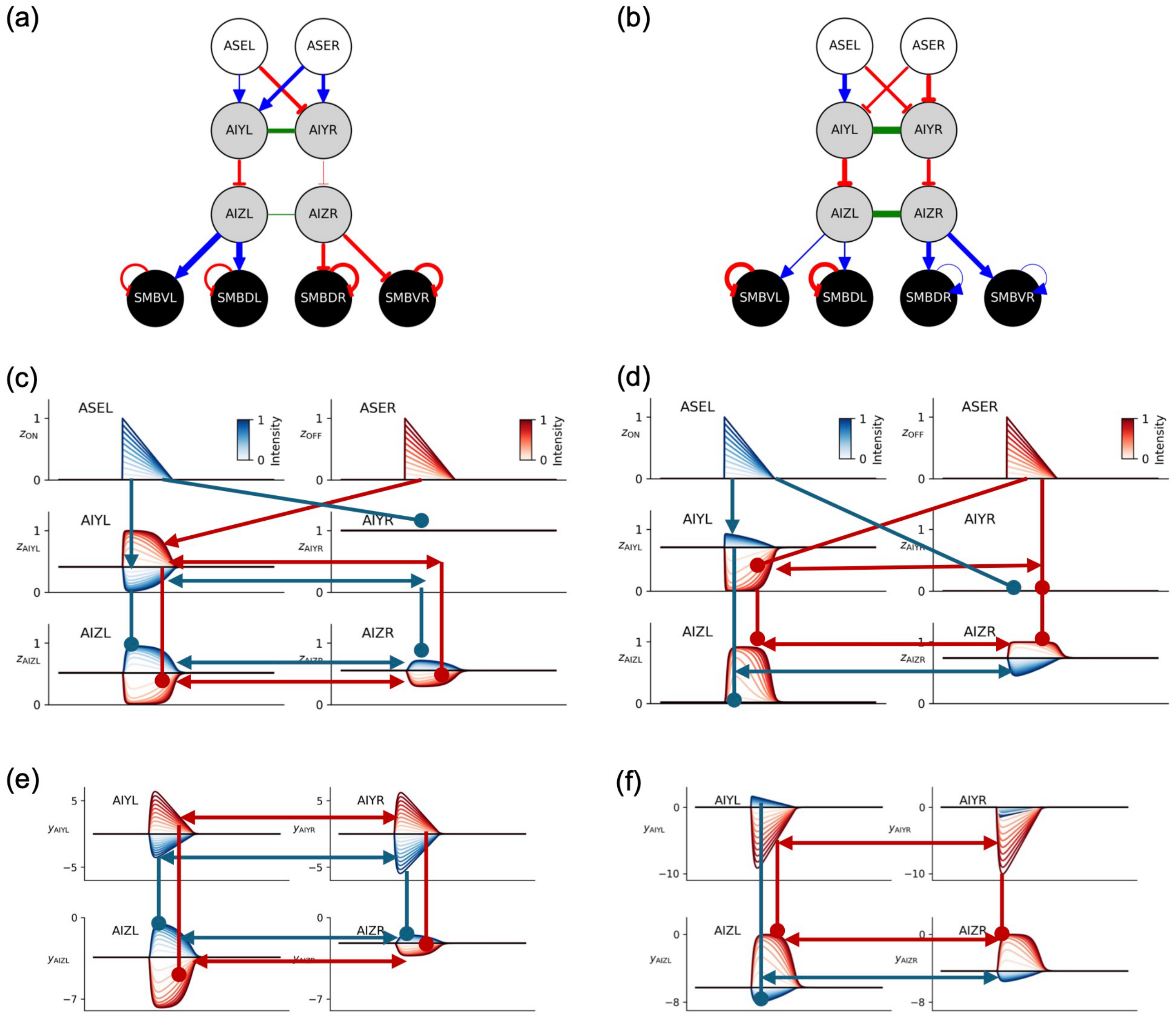


**Figure S1. Signal transmission through chemical synaptic connections and electrical gap junctions.** (a) and (b) The most optimized neural circuits developed without (a) and with (b) the constraints mentioned in the text. For purposes of reference, the same as in Fig. 3a and 3b are shown again. (c) and (d) The neurotransmitter release, $z_i$, from each neuron in the network (a) and (b) in response to step changes in salt concentration of positive (blue) and negative (red) is shown in (c) and (d), respectively, along with the signaling through the chemical synaptic connection and the electrical gap junction. The $z_i$s shown herein are the same as those shown in Fig. 3c and 3d. (e) and (f) The membrane potential $y_i$ that was observed in the network (a) and (b) is shown in (e) and (f), respectively. In (c)-(f), the blue (red) arrow and the blue (red) line with an ending in a filled circle show the excitatory and inhibitory synaptic transmission, respectively, in response to positive (negative) step changes in salt concentration. The blue (red) bidirectional arrow indicates gap junction signal transmission, in response to positive (negative) step changes in salt concentration.

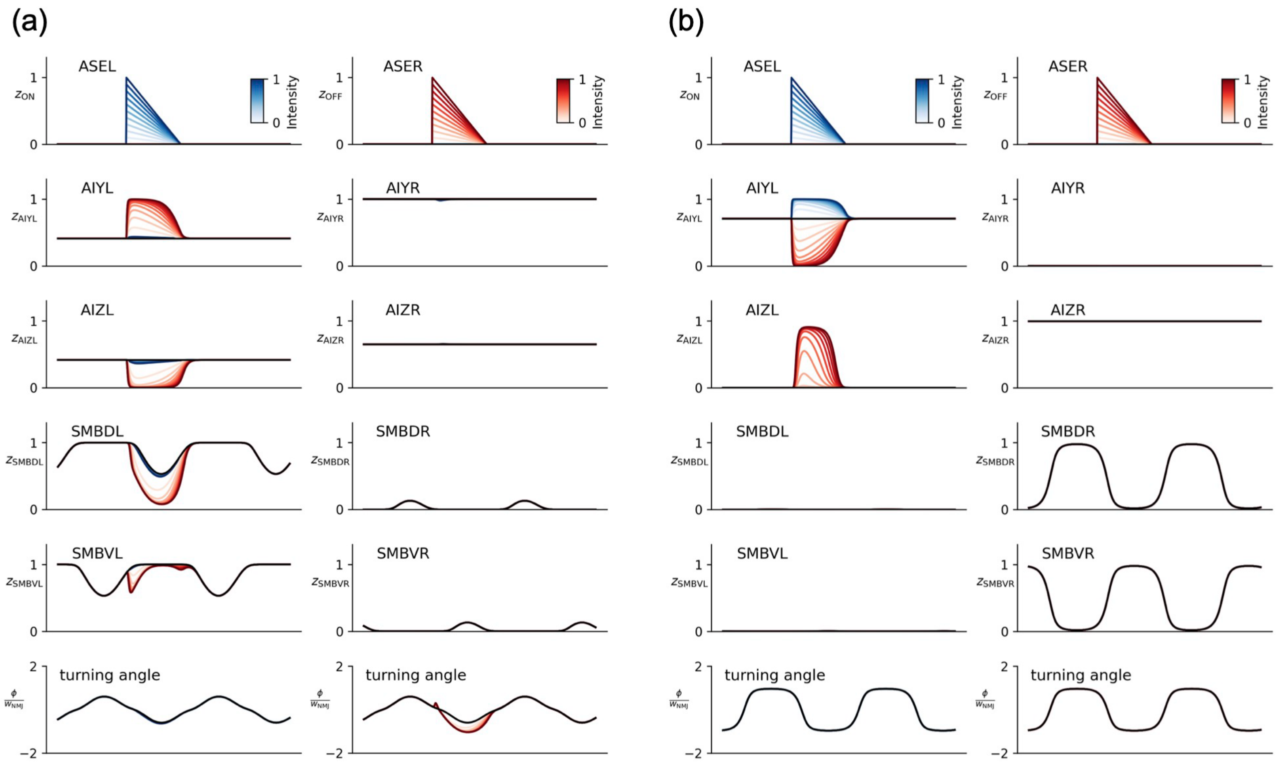


**Figure S2. Blocking electrical gap junctions significantly has a marked effect on neurotransmitter release $z_i$ and the resulting turning angle $\varphi$.** (a) and (b) The most optimized neural circuits without (a) and with (b) the constraints. With the exception of the gap junctions that have been blocked, all other parameters remain consistent with those illustrated in Fig. 3.

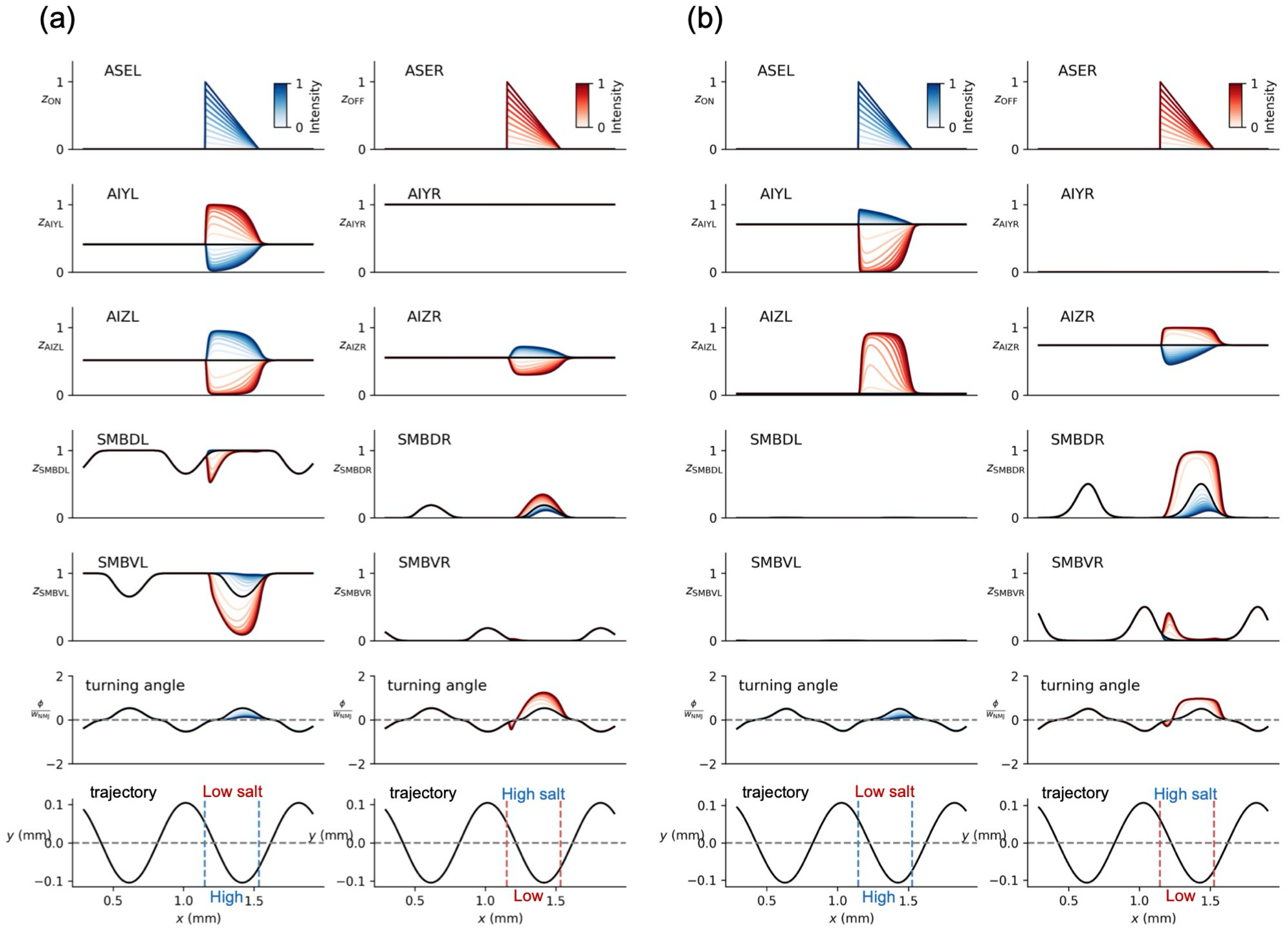


**Figure S3. The changes in the turning angles $\varphi$ in response to step changes in the salt concentration of positive and negative at the time half a cycle later than those shown in Fig. 3.** (a) and (b) The results obtained from the most optimized neural circuits without and with the constraints are shown in (a) and (b), respectively. With the exception of the timing of the step changes in salt concentration, all other conditions are identical to those depicted in Fig. 3.

The most optimized model with the constraints (also shown in Fig. 5a).

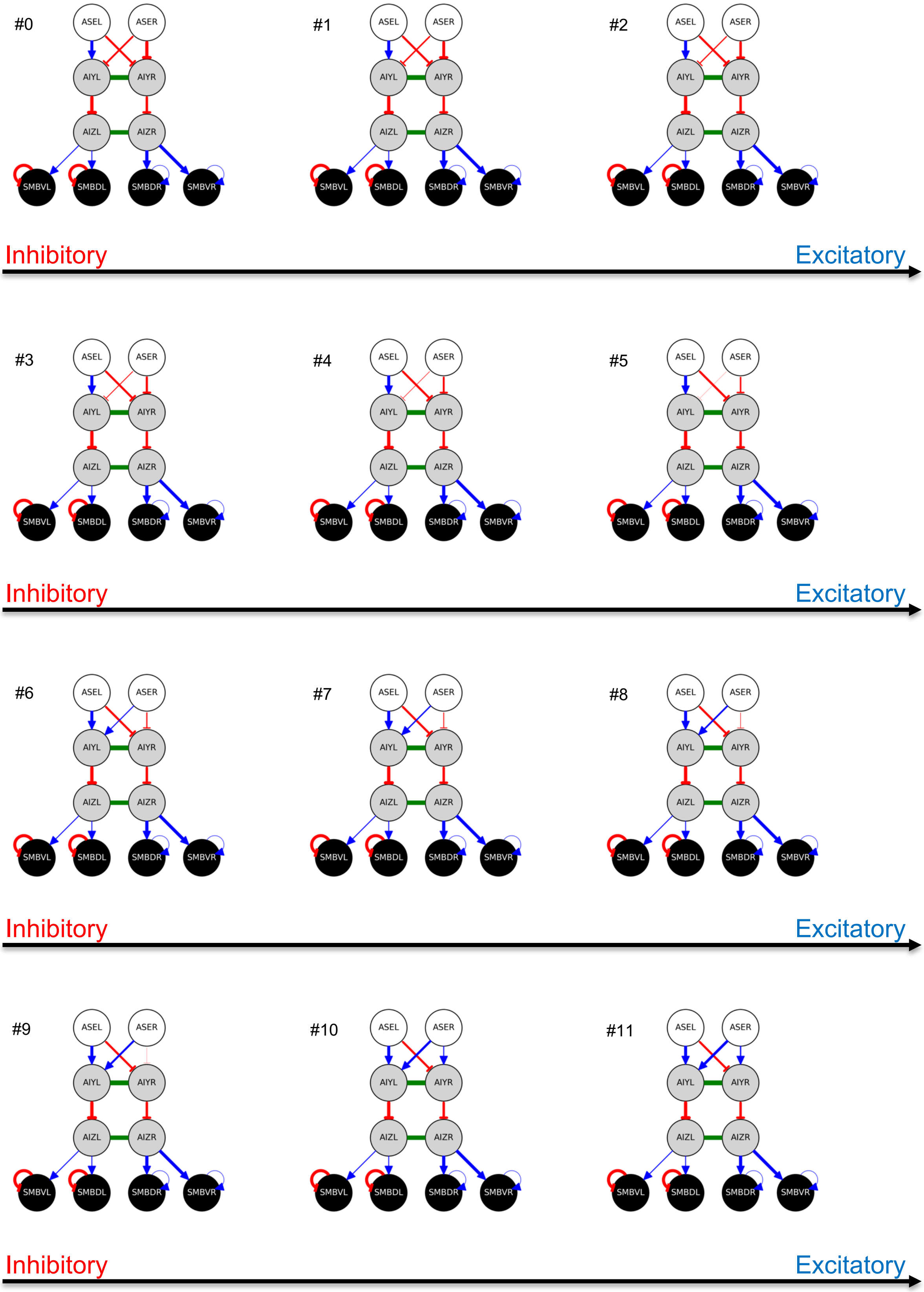

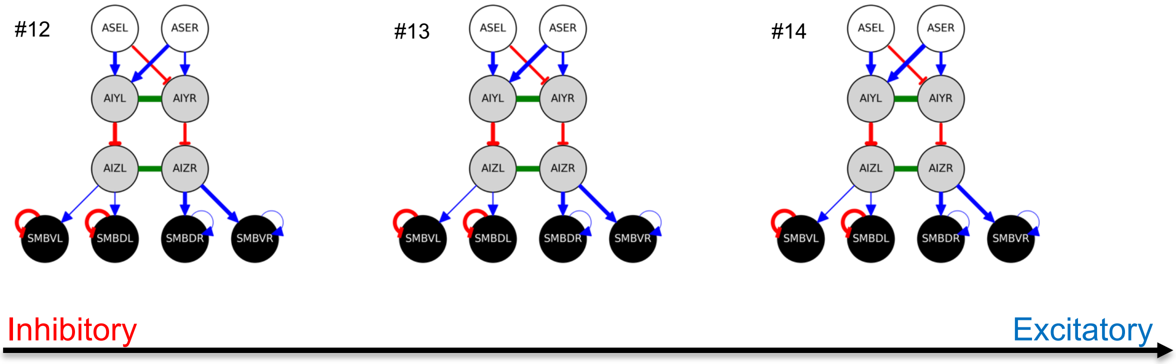


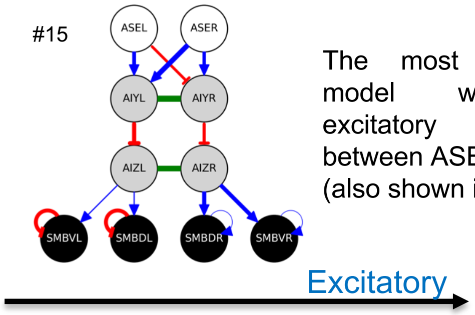


The most optimized model with the excitatory connection between ASER and AIY (also shown in Fig. 5b).

**Figure S4. The weight of the ASER-AIY synaptic connection, $w_{ji}$, in the most optimized network with the constraints is increased from a negative value (inhibitory connection) to a positive value (excitatory connection) by introducing an increment of 1.5 to $w_{ji}$ at each step.**

The most optimized model with the constraints, which is considered to correspond to a well-fed individual cultivated in an environment with a higher salt concentration than the current one (as also indicated by the black symbols in Fig. 5c).

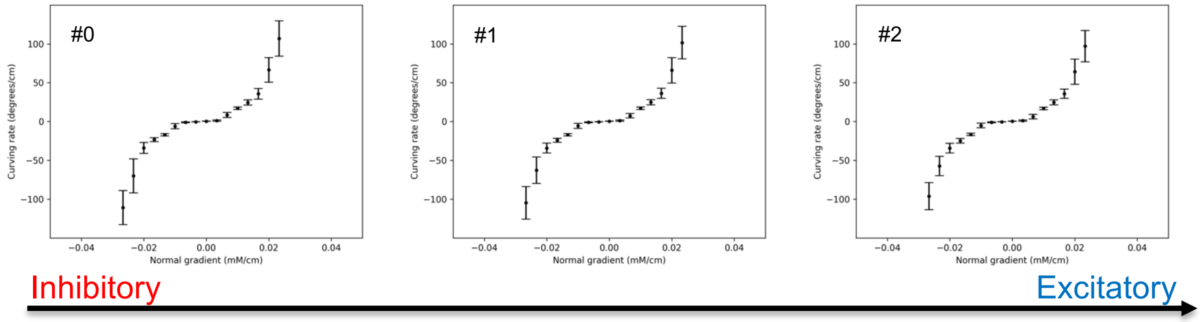


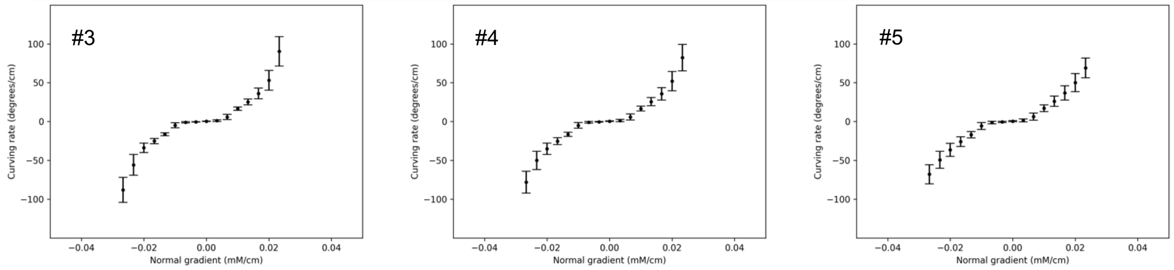


Inhibitory Excitatory

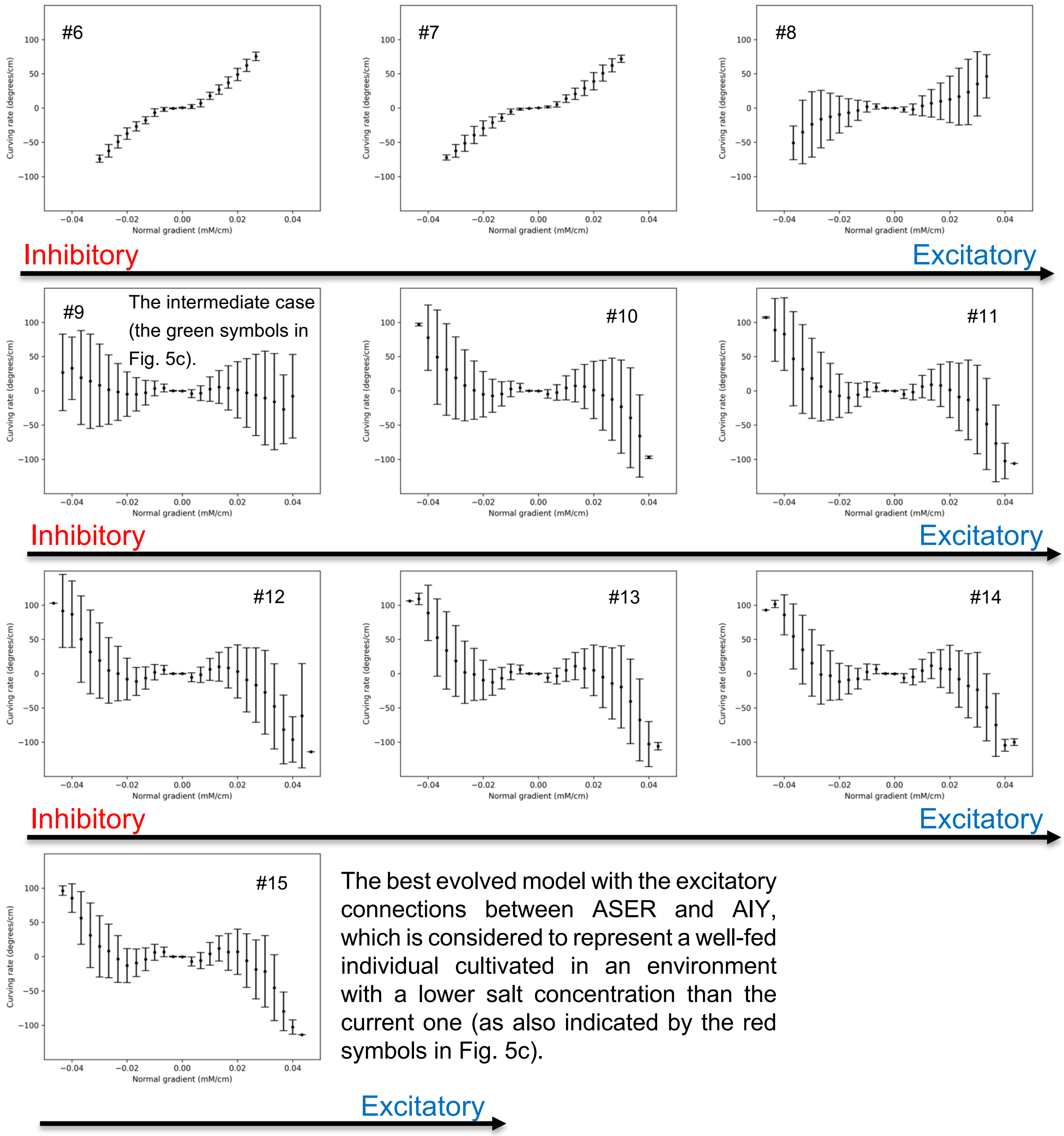


**Figure S5. As the weight of the ASER-AIY synaptic connection $w_{ji}$ is increased from negative (inhibitory connection) to positive (excitatory connection) in the most optimized network with the constraints by introducing an increment of 1.5 to $w_{ji}$ at each step, the curving rate is varied from an increasing function with an increasing normal gradient of salt concentration to a function that shows a decreasing trend.** The curving rate shown in the #0, #9, and #15 is also shown as that for the inhibitory, intermediate, and excitatory connections, respectively, in Fig. 5c.

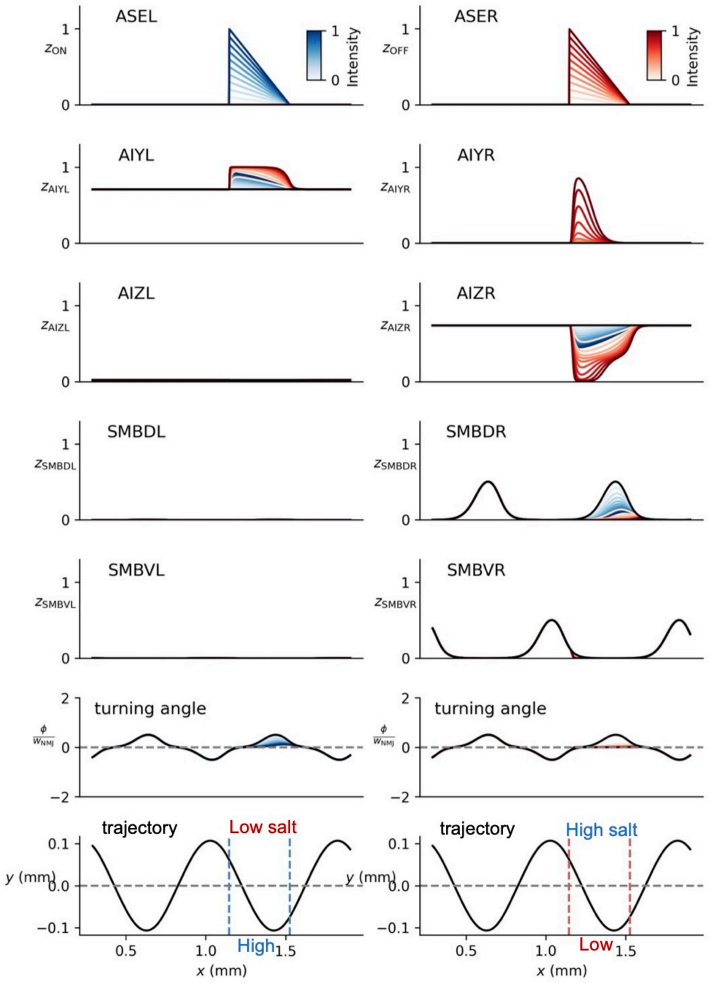


**Figure S6. The changes in the turning angles $\varphi$ in response to step changes in the salt concentration of positive and negative at the timing half a cycle later than those shown in Fig. 5e.** The changes in $\varphi$ in response to step increases in $z_{ON}$ and $z_{OFF}$ at the timing of half a cycle later ensured that the regulation mechanism of $\varphi$ discussed in the text (Fig. 5e) was maintained.

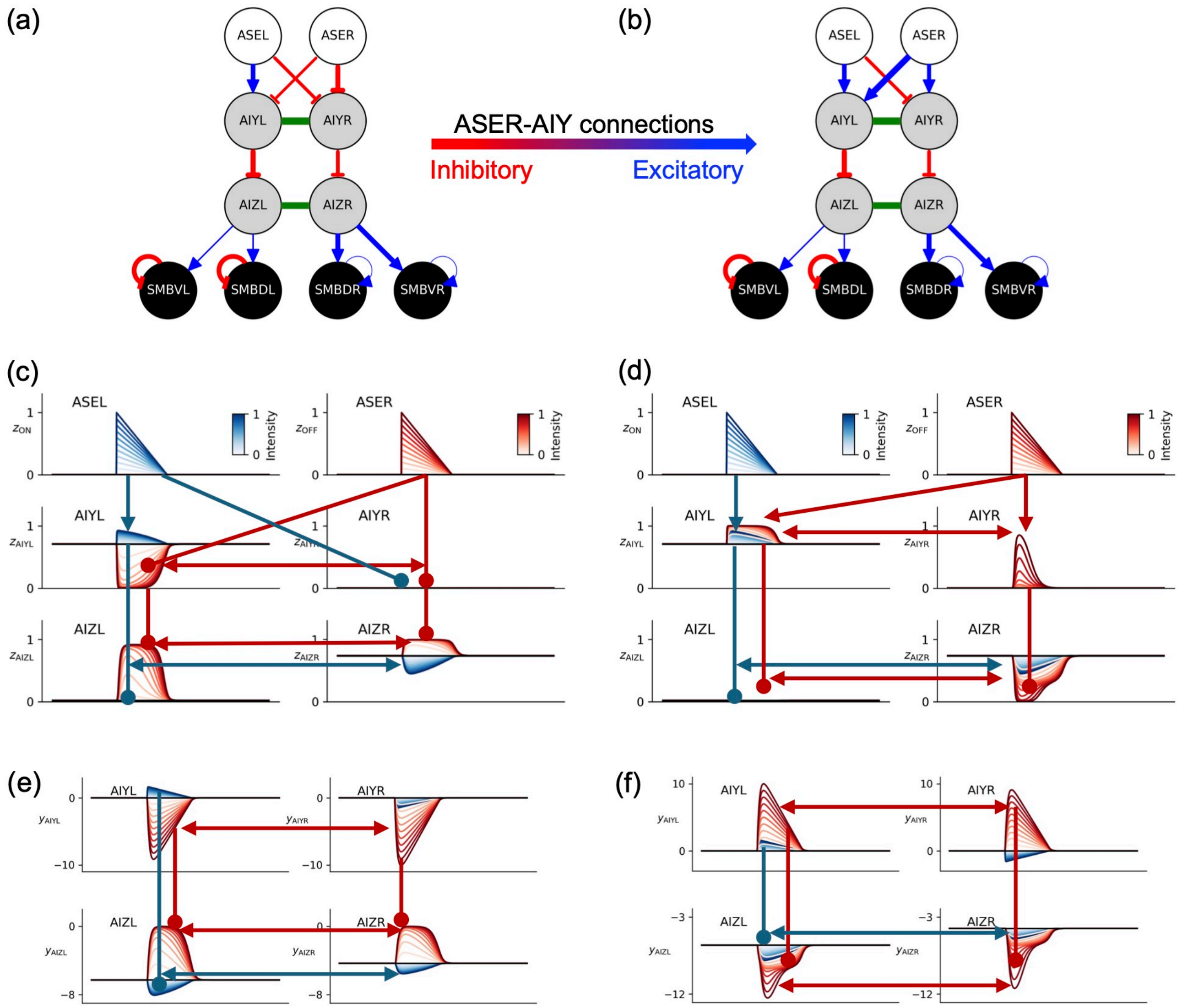


**Figure S7. Signal transmission through chemical synaptic connections and electrical gap junctions.** (a) and (b) For purposes of reference, those shown in Fig. 5a and 5b are presented again in (a) and (b), respectively. (c) The neurotransmitter release $z_i$ in response to step changes in the salt concentration of positive (blue) and negative (red) for the most optimized circuit with the constraints shown in (a), is shown together with the signal transmissions through the chemical synaptic connections and electrical gap junctions. (d) The same as (c) is shown, except that the inhibitory connections between the ASER and AIY are replaced with the excitatory connections, as shown in (b). The $z_i$s shown in (c) and (d) are the same as those shown in Fig. 3d and 5e, respectively. (e) and (f) The membrane potentials $y_i$ resulting in $z_i$, as illustrated in (c) and (d), are shown in (e) and (d), respectively. In (c)-(f), the blue (red) arrow and the blue (red) line with an ending in a filled circle indicate the excitatory and inhibitory synaptic connections, respectively, in response to step changes in the salt concentration of positive (negative). The blue (red) bidirectional arrow indicates the transmission via gap junctions, in response to positive (negative) step changes in salt concentration.

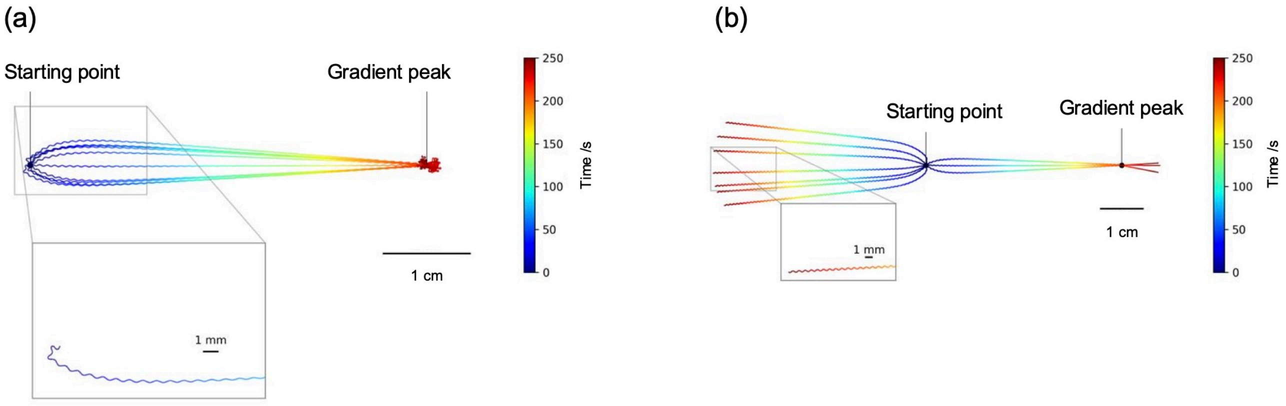


**Figure S8. The trajectories of the worm's locomotion simulated by the network shown in Fig. 5a and 5b.** (a) For purposes of comparison, the same trajectories depicted in Fig. 2b are presented, again. (b) The trajectories given by the network depicted in Fig. 5b, wherein the inhibitory connections between the ASER and AIY in the most optimized model with the constraints have been substituted with excitatory connections. The majority of the trajectories exhibited a directionality that was opposite to the peak of the salt concentration gradient. However, a subset showed a trajectory where the worm proceeded towards the peak of the gradient and passed it without any discernible response.

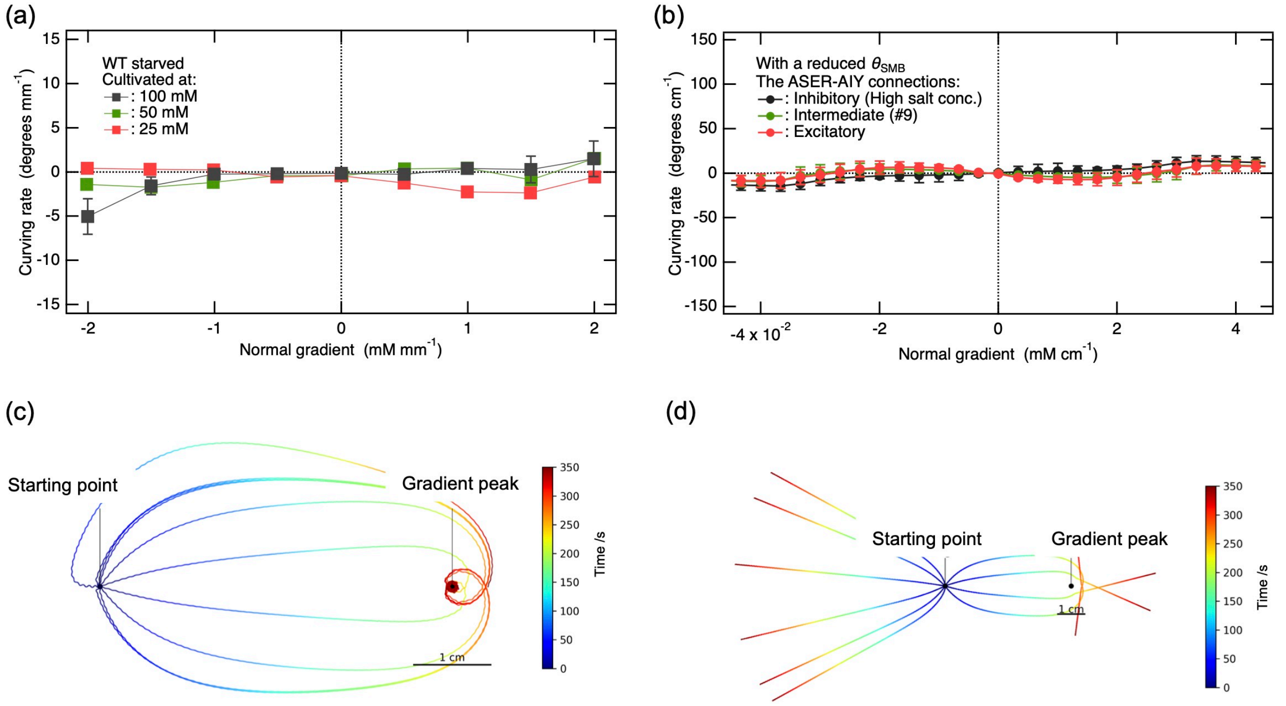


**Figure S9. The inhibition of SMB activity by reducing the bias term $\theta_{\text{SMB}}$ has the effect of suppressing the salt memory-dependent preference behavior observed in klinotaxis, similar to that illustrated in Fig. 6b.** (a) For purposes of comparison, the same experimental data on the curving rate as presented in Fig. 6a is provided here again. (b), (c) and (d). The figures shown in (b), (c) and (d) are the same as shown in Fig. 6b, 6c, and 6d, respectively, except that in the most optimized models depicted in Fig. 5c, the bias terms for the SMB motor neurons were reduced, rather than the synaptic connections between AIZ and SMB being weakened as was done in Fig. 6.
